# Identification of selection and inhibition components in a Go/NoGo task from EEG spectra using a machine learning model

**DOI:** 10.1101/705525

**Authors:** Bambi L. DeLaRosa, Jeffrey S. Spence, Michael A. Motes, Wing To, Sven Vanneste, John Hart, Michael A. Kraut

## Abstract

Prior Go/NoGo studies have localized specific regions and EEG spectra for which traditional approaches have distinguished between Go and NoGo conditions. A more detailed characterization of the spatial distribution and timing of the synchronization of frequency bands would contribute substantially to the clarification of neural mechanisms that underlie performance of the Go/NoGo task. The present study used a machine learning approach to learn the features that distinguish between ERSPs involved in selection and inhibition in a Go/NoGo task. A neural network classifier was used to predict task conditions for each subject to characterize ERSPs associated with Go versus NoGo trials. The final model accurately identified individual task conditions at an overall rate of 92%, estimated by 5-fold cross-validation. The detailed accounting of EEG time-frequency patterns localized to brain sources (i.e., thalamus, preSMA, orbitofrontal cortex, and superior parietal cortex) provides elaboration on previous findings from fMRI and EEG studies and more information about EEG power changes in multiple frequency bands (i.e., primarily theta power increase, alpha decreases, and beta increases and decreases) within these regions underlying the selection and inhibition processes engaged in the Go and NoGo trials. This extends previous findings, providing more information about neural mechanisms underlying selection and inhibition processes engaged in the Go and NoGo trials, respectively, and may offer insight into therapeutic uses of neuromodulation in neural dysfunction.

## 1 Introduction

Go/NoGo tasks have been used in cognitive neuroscience to explore brain mechanisms underlying inhibitory control, selection, and, more broadly, cognitive control (e.g., see [1]). Cognitive control deficits and inhibitory dysfunction, in particular, have been linked to clinical conditions such as Attention Deficit Disorder (ADD) [2], traumatic brain injury (TBI) ([3], and schizophrenia [4], and a host of studies have sought to understand the neural mechanisms supporting inhibition and selection. Go/NoGo tasks require responding to designated “Go” stimuli (e.g., a green square) and withholding responding to other designated “NoGo” stimuli (e.g., a red square). The proportion of Go stimuli is typically higher than NoGo stimuli (e.g., 80% Go and 20% NoGo) to establish a biased expectation to respond on any given trial. Go/NoGo task performance then depends on the abilities both to select to respond to Go stimuli as well as inhibit the established prepotent response to NoGo stimuli.

Scalp-recorded EEG signals consist of mixed signals from various neural sources. The preponderance of electroencephalography (EEG) research on selection and inhibition in Go/NoGo paradigms has focused on N2 and P3 ERP components. On both Go- and NoGo trials, both a negative deflection of the ERP occurring between 250 and 350 ms (i.e., N2), and a positive deflection occurring between 300 and 600 ms (i.e., P3) have been observed, both with a midline frontal to parietal distributions and with NoGo waveform amplitudes exceeding that of Go [5, 6, 7, 8]. The N2-P3 components have been shown to relate to inhibitory processes, but the relationships of these components to specific inhibitory functions remain to be fully specified [9].

In studies examining EEG oscillatory dynamics in Go/NoGo tasks, frontal potential fluctuations in the theta range (i.e., 4-8 Hz) have been have been posited to index processes related to selection and inhibition [10, 11], with theta-power accentuated in inhibition compared to selection. In a previous study performed by our group, we found that theta-power between 200 and 600 ms after the onset of a trial was greater during NoGo than during Go trials and found local maxima for the theta-power increase over two fronto-centeral regions corresponding to frontal pole and pre-SMA, with theta oscillations being coherent between the two regions for the NoGo condition [12]. Furthermore, EEG source localization analysis of Go/NoGo data has suggested anterior-posterior distinctions between NoGo and Go signal generators, with NoGo signal generators having a more anterior localization [13]. Magneto-encephalography (MEG) findings also suggest localization of inhibitory control signals to fronto-central regions [14], and midline theta changes in EEG correspond to medial-frontal/anterior cingulate theta fluctuations in MEG [15].

Functional MRI studies have provided anatomic localization in Go/NoGo tasks. Functional MRI signal from the pre-SMA was increased for both Go and NoGo trials [16], suggesting involvement in both response selection and inhibition. Additionally, the preSMA showed greater activation on NoGo than on Go trials, consistent with the time-frequency data, suggesting additional involvement in response inhibition. Inhibition-related effects (i.e., greater fMRI signal change on NoGo compared to Go trials) were observed within right tempro-parietal, right inferior and middle frontal, middle temporal, and pre- and post central gyrus regions. Furthermore, in a meta-analysis of fMRI studies of response inhibition, NoGo stimuli preferentially were found to engage a network including right dorsolateral prefrontal cortex (DLPFC), right inferior frontal gyrus, right inferior parietal lobule (IPL), preSMA and anterior cingulate cortex (ACC), and the insula [1].

Lesion studies also have helped delineate essential roles for the frontal pole and the preSMA in selection/inhibition tasks. Lesions of the frontal pole have been shown to increase reaction times substantially in Go/NoGo tasks [17], and lesions of pre-SMA areas have been shown to increase the number of false-positives in the NoGo condition [18, 17]. These lesion-related findings support the concept that compared to more rostral brain structures, one role the pre-SMA area plays is in inhibiting the motor action.

While there is an accumulating body of data regarding the spatial distribution of frequency-band specific EEG changes and localization of function via fMRI and lesion studies, there has been less attention paid thus far as to the temporal sequence of these changes. A combination of the spatial and the temporal attributes of the EEG change would contribute substantially to the clarification of neural mechanisms that underlie performance of the Go/NoGo task, and response inhibition more generally. In the present study, we have investigated a Go-NoGo task while recording EEG and evaluating event-related spectral perturbations (ERSP) [19] of independent EEG components. We utilized a neural network classifier to predict, with maximal accuracy, the Go and NoGo conditions from the component signals for each subject. Our approach is thus a machine learning approach to identify the spectral, temporal, and spatial electrophysiological features involved during selection and inhibition. The features learned by the neural network model included the identification of the sources of the components, the multiple frequency content of components, and the timing of maximal synchronization or desynchronization of activity that differentiate Go and NoGo trials.

## 2 METHODS AND MATERIALS

### 2.1 Subjects

Fifty-nine subjects (31 M, 28 F) were recruited from the University of Texas at Dallas community via word of mouth and web-based advertising. All subjects were between the ages of 18 and 35. The mean age of the group was 23.5 (SD 4.1). There were 3 African American, 1 American Indian, 16 Asian, 24 Caucasian, 10 Hispanic, and 5 Multiracial participants. The subjects were all college students or graduates with at least 12 years of education.

Subjects were screened, per exclusion criteria, to be free from a history of traumatic brain injury and other significant neurological issues (stroke, seizure disorders, history of high fevers, tumors, or learning disabilities). Exclusion criteria also included left-handedness, use of alcohol or other controlled substances within 24 h of EEG administration, and medications other than over-the-counter analgesics and oral contraceptives. Two subjects were excluded from the analysis due to corrupted EEG files.

Informed written consent was collected from each subject according to the rules of the Institutional Review Board of The University of Texas at Dallas. This study was conducted according to the Good Clinical Practice Guidelines, the Declaration of Helsinki, and the US Code of Federal Regulations.

#### 2.1.1 Stimuli

We used a basic Go-NoGo paradigm consisting of 160 (80%) ‘Go’ stimuli (a drawing of a specific car) for which the subject was to press a button and 40 (20%) ‘NoGo’ stimuli (a drawing of a specific dog) for which the subject was instructed to withhold a response. The stimuli were presented for 300 ms followed by a fixation point (+) for 1700 ms. All of the stimuli were black line drawings fitted to a white 600 × 600-pixel square. The instructions were to press the button for all cars but to do not press the button for anything else. Instructions were given verbally and displayed on the computer screen prior to the task.

### 2.2 EEG Recording

Continuous EEG was recorded from a 64-electrode Neuroscan Quickcap using Neuroscan SynAmps2 amplifiers and Scan 4.3.2 software, with a reference electrode located near the calvarial vertex. Data were sampled at 1 kHz, with electrode impedances typically below 10 kΩ. Additionally, bipolar electrooculographic data were recorded from two electrodes to monitor blinks and eye movements (positioned vertically at the supraorbital ridge and lower outer canthus of the left eye). The continuous EEG data were offline high-pass filtered at 1 Hz and low-pass filtered at 50 Hz using a finite impulse response (FIR) filter.

### 2.3 EEG Pre-Processing

We analyzed the EEG data using scripts developed in our lab that implement functions from EEGLAB version 13.1 [20]running under Matlab 7.11.0. Preprocessing consisted of down-sampling to 512 Hz, removing data recorded from poorly functioning electrodes, and correcting for stereotyped artifacts including eye blinks, lateral eye movements, muscle, line noise, and heart rate using the runica algorithm [20, 21], an implementation of the logistic infomax independent component analysis algorithm of Bell and Sejnowski [22]. Stereotyped artifacts were identified by visual inspection of the spatial and temporal representation of the independent components. Continuous data were then segmented into 3-second non-overlapping epochs spanning from 1000 ms before to 2000 ms after the presentation of the visual stimuli. Epochs containing high amplitude, high frequency muscle noise, and other irregular artifacts were removed retaining on average 85 percent of all epochs. Finally, missing electrodes were interpolated and data were re-referenced to the average reference [23].

### 2.4 Obtaining Independent Sources of EEG Signals

The cleaned and filtered EEG data were reshaped for purposes of isolating independent sources through Independent Components Analysis (ICA). For each subject and at each channel, the task condition trials were mean centered and concatenated. Each subject’s concatenated trials were then scaled to have equal variance. At this stage all subjects were concatenated to yield a large EEG data frame containing the set of 3-second trials (−1 to 2 seconds post-stimulus), down-sampled at 128 Hz, across task conditions and subjects for each channel. We obtained the components from the ICA using the extended infomax option in EEGLAB’s runica function and temporarily kept all components for purposes of fitting single dipole scalp projections.

Taking advantage of the fact that independent components are more dipole-like than the raw channel-level EEG signals [24], we used EEGLAB’s dipfit function (boundary element method) to localize equivalent dipole sources of independent component scalp maps. This procedure allowed us to obtain approximate locations within the brain of the independent component sources themselves. We discarded components that were not isolated to gray matter, and we discarded components whose residual variance was greater than 15% from the fit of the independent component scalp maps to the scalp projections of single equivalent dipoles. The process of attrition left 22 viable independent components and the approximate locations within the brain of their equivalent dipole sources.

### 2.5 Spectral Estimation of Independent Components

Before estimation of the frequency spectra of the derived components from ICA, we isolated the subject-level data by partitioning each subject’s respective segments of the components for both task conditions. In addition, we saved the ICA weights for purposes of projecting individual subjects and their task conditions into the derived component space. This latter step was important for cross-validation (see details below) in which subjects who were left out of the ICA and subsequent dipole fitting stage could be projected into the appropriate space for later prediction.

Once subject-level component segments for each condition were isolated, we calculated the frequency spectrum using Matlab’s newtimef function in cosine tapered windows across .05-second segments of the trial window. The modulus of the Fourier transform was converted to log scale, and each post-stimulus trial segment was baseline-corrected using the .75 seconds pre-stimulus onset period as baseline. The frequency range was from 1 Hz to 30 Hz with a frequency resolution of 1 Hz, and the temporal range was from −.75 seconds to 1.75 seconds post-stimulus onset with an effective temporal resolution of .05 seconds. Finally, the time-frequency spectra for each task condition was trial averaged for each subject.

At this stage of data processing the subject-level arrays for each task condition consisted of log spectra for 22 independent components localized by fitting projections of equivalent dipole sources, and a time-frequency panel covering up to 30 Hz for a 2.5-second time window around the time of stimulus onset.

### 2.6 Overview of the Machine Learning Approach

Data features, denoted by *X*, were inputs to the prediction model, trained to learn the important elements that could classify the task condition. *X* is a *p*-length vector comprising scaled, log power spectra at localized sources of independent EEG components, average frequency intervals, and .8-second time windows beginning at the stimulus onset for each condition. The output of the prediction model takes values in the set *C* = {*Go, NoGo*}, which are the task conditions. We proceeded to train a model that would estimate prediction functions *f*_*k*_(*X*), *k* ∈ *C*, which are condition probabilities given the input features *X*. A single subject’s experimental condition was then predicted as *Ĉ*(*X*) = argmax_*k*_ *f*_*k*_(*X*) the most probable class, where “^” denotes the estimate derived from the trained prediction model. The final prediction model included the appropriate *p*-length vector *X*, as well as all model parameters (see details below), that minimized an estimate of test error, defined as the proportion of *C*_*k*_(*X*) misclassified as *C*_ℓ_(*X*). Model choice and the estimate of its final test error were both obtained through 5-fold cross-validation (CV).

### 2.7 Deriving Input Features X for the Prediction Model

We chose a single-layer neural network learning model, with *m* = 1, …,*M* derived units in the “hidden” layer, to classify experimental conditions for single subjects. This model has *M* (*p* + 2) + 2 parameters to estimate, with *X* having *j* = 1, …, *p* features. Since this model is overparameterized, one of the additional parameters is an *L*__2__ penalty to impose constraints on the other *M*(*p* + 2) + 1 and to prevent overfitting (details below).

To alleviate a significant computational burden, but more importantly for parsimony and interpretive value, we reduced p to a number beyond which we no longer reduced cross validation (CV) error. This was accomplished in two ways—fixing the values of time and frequency and finding a suitable subset of components. First, we fixed the time window to be only 0.8 seconds following stimulus onset, and we averaged frequencies within the intervals 4-8 Hz, 9-10 Hz, 11-12 Hz, 13-20 Hz, and 21-30 Hz. These intervals roughly correspond to theta band, lower and upper alpha band, lower and upper beta band, respectively. Secondly, we ran combinations of components through CV to obtain a subset with minimum CV error. Specifically, we fixed two which we knew from prior literature and experience to be important (components 1 and 5, see Table 1), and we proceeded to add groups of 3-5 components at a time, followed by reductions of 1-2. This process of forward and backward selection, although not nearly exhaustive, relatively quickly allowed us to find a subset with minimum CV error. Adding any single component to this subset or removing any single component from this subset only increased CV error.

**Table 1:**
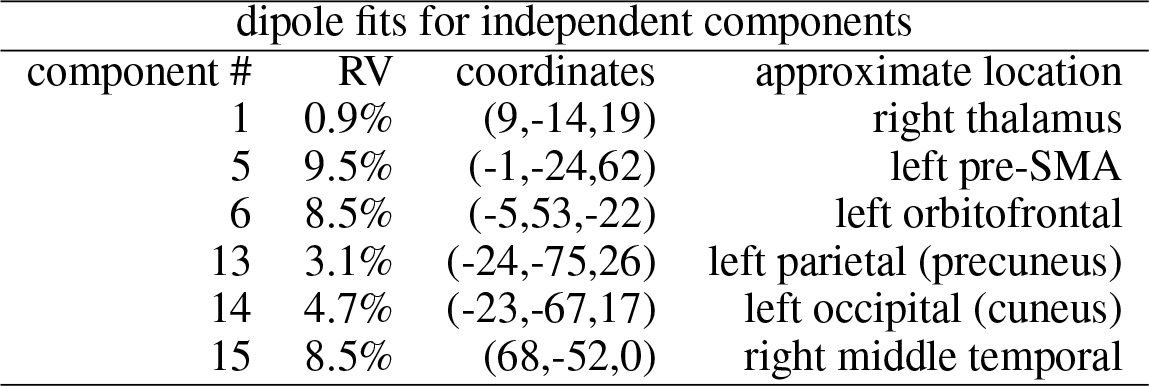
Component numbers from the ICA and their approximate locations based on best fits of the component scalp maps to the scalp projections of single equivalent dipoles. The assessment of each fit is given by % residual variance (RV).

Inputs *X* to the neural network model require scaling of the features. We accomplished this by centering each subject and component and setting their respective variances equal to 1. Additionally, we adjusted temporally based on reaction times to Go trials. That is, we regressed time to peak (in absolute value) on average reaction times and adjusted time-frequency windows accordingly. We applied the same adjustment to NoGo trials, which implicitly assumes that Go and NoGo reaction times are approximately proportional. This adjustment was fairly minimal but it significantly reduced CV error.

### 2.8 Model Selection

Choosing the final neural network prediction model required finding the number, *M*, of derived units in the “hidden” layer, the subset of independent components making up part of the *p*-length vector *X*, *L*_2_ penalty λ, and the model parameter set *θ* = {*α*′, *β*′} which minimizes the penalized cross-entropy loss function

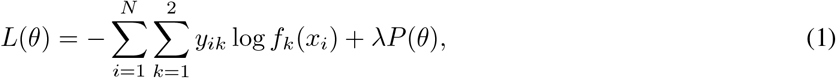

where *y*_*ik*_ is an indicator of task condition, 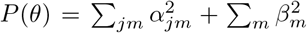 is a penalty functional, and λ is the *L*_2_ parameter (i.e., weight decay parameter) controlling the influence of the penalty when minimizing the loss function, *L*(*θ*). The greater λ is, the more constrained are the size of the parameters *α*′ and *β*′ that are implicitly contained in the prediction function *f*_*k*_(*x*_*i*_). As noted above, λ is an important consideration due to the fact that the model is overparameterized.

Another important aspect of λ is its influence on the models’ robustness to initialization parameters. Minimizing *L*(*θ*) is a nonlinear optimization and requires starting values for initialization of the BFGS quasi-Newton method. If the CV error changes minimally for different sets of initialization parameters, we call the model “robust” to those starting values, and λ has a direct influence on that robustness property (see Figure 1 in Results). For all iterations of model selection we chose random uniform initialization parameters between .7 and .7.

**Figure 1:**
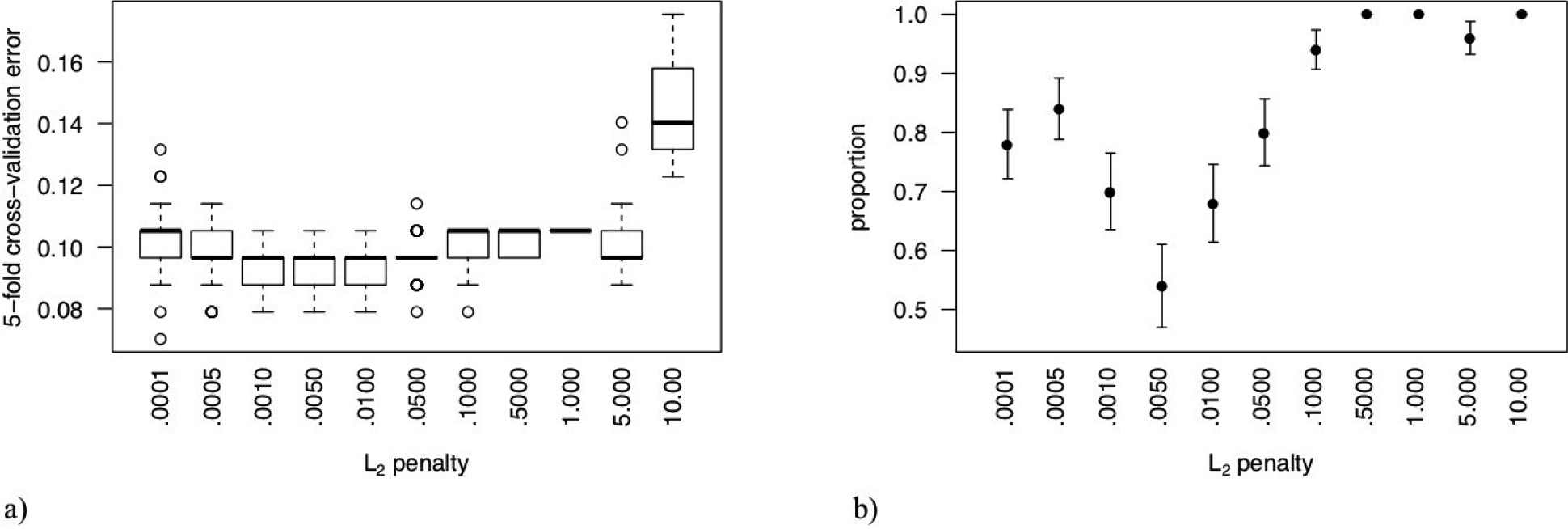
a) Estimates of the proportion of *C*_*k*_(*X*) misclassified as *C*_ℓ_(*X*) by 5-fold cross-validation (CV) as a function of the *L*_2_ penalty. Each boxplot is a distribution summary of error estimates from 50 sets of random uniform initialization parameters in the neural network based on *M* = 4 derived units in the single layer. b) Proportion of the 50 initialization sets that yield a final CV prediction error greater than 0.09. An *L*_2_ penalty equal to .005 yields models that are most robust to initial starting values and have a higher proportion of CV prediction error rates equal to that of our final model (.079).

The issues noted here were implemented as various options in the nnet function of the nnet package for the R statistical computing environment (http://r-project.org). All parameters and features - *M*, *θ* = {*α*′, *β*′}, λ, and *p*- were chosen to minimize CV error.

### 2.9 5-Fold Cross-Validation

Neural network learning models are highly adaptive to training data and will overfit such that predictions using independent input features *X* will generally be poor. We implemented 5-fold cross-validation (CV) to find an optimal model that will generalize well. As noted, CV was used both to find the best model and to estimate its test error for generalizability. We emphasize that all aspects of the data processing and modeling stream — from ICA and dipole fitting to the search for feature/parameter sets {*M*, *θ*, λ, *p*} — were included in the CV.

For this procedure we randomly partitioned the total sample of subject/conditions, 2 × 57 = 114, into roughly 5 equal parts. For the *k*^*th*^ partition, or fold comprising 20% of the data, we defined a neural network learning model for a particular {*M*, *θ*, λ, *p*}-set, trained on the other 4 parts comprising 80% of the data, then calculated misclassification error (i.e., test error) for the *k*^*th*^ fold using the model built without it. This process was repeated five times, once for each of the *k* = 1, …, 5 folds; and the cross-validation error was estimated by combining the 5 individual estimates. A summary of steps for each candidate model was the following:

1. Divide total sample into 5 folds at random
2. Obtain independent components through ICA using all of the samples *except* those in fold *k*
3. Fit equivalent dipole models to component scalp maps using all of the samples *except* those in fold *k* (Note: component numbers generally did not come out in the same order across the folds, but they were easily matched by finding the closest dipole location within 5 mm)
4. Choose the p-length vector X, particularly a subset of independent components, using all of the samples except those in fold k
5. Train the neural network learning model for each {*M*, *θ*, λ, *p*} set using all of the samples except those in fold *k*
6. From the trained model obtain the classification *Ĉ*(*X*) and its misclassification error rate for the samples in the hold-out fold *k*
7. Accumulate errors from all 5 folds to produce the cross-validation estimate of test error for the candidate model and its associated {*M*, *θ*, λ, *p*}-set.

## 3 RESULTS

### 3.1 Final Neural Network Learning Model

At the end of model selection we isolated *p* = 240 features comprising input vector *X*: 6 components × 5 frequency intervals × 8 time units between 0 and .8 seconds post-stimulus. The frequency intervals were, approximately, theta band, lower and upper alpha band, lower and upper beta band; and the six components are given in Table 1. Approximate locations of these sources are shown, as well as the coordinates of the equivalent dipole having a scalp projection that matches closely with the given component’s scalp map.

Cross-validation chose *M* = 4 units in the hidden layer of the neural network and λ = .005. Figure 1 (a) shows CV curves as a function of the *L*_2_ penalty for models with *M* = 4 derived units and *p* = 240 features in *X*. The box plots are the CV error distribution summaries of 50 different initialization parameter sets for each value of λ. Although the CV error summaries seem relatively flat near λ = .005, Figure 1 (b) shows that λ = .005 yields models that are more robust to initialization parameters (i.e., lowest proportion of initialization parameter sets with CV error above .09).

The final learning model has very good accuracy in predicting task conditions at the subject level. Although the model can predict the conditions from the training data with 100% accuracy, the CV error is a better reflection of generalization error when predicting task conditions from independent subjects. These rates are shown in the confusion matrix given in Table 2. The off-diagonal entries show the CV error rates for each task condition with an overall error rate of .079 (standard error = .016).

**Table 2:**
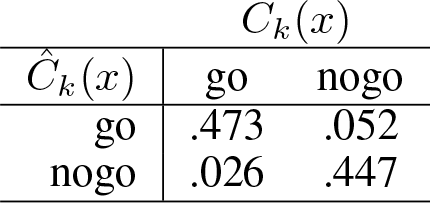
Table 2. Confusion matrix, estimated by 5-fold cross-validation, for the neural network learning model of task condition. *Ĉ*_*k*_(*x*) is the predicted condition from the model, and *C*_*k*_(*x*) is the true experimental condition. The overall test error rate is .079 (se = .016).

| $\hat{C}_k(x)$ | $C_k(x)$ | |
| --- | --- | --- |
|  | go | nogo |
| go | .473 | .052 |
| nogo | .026 | .447 |

### 3.2 Interpretation of the Prediction Model

Interpretation of the features in *X* that are essential for accurate task condition predictions requires a more detailed look into the units of the hidden layer of the neural network. These derived units contain the initial information from *X* that contributes to the prediction functions *f*_*k*_(*X*). For each of the *m* = 1, …, 4 units we used the corresponding model parameters in *α′*, which are the subset of parameters that relate *X* to each individual unit of the hidden layer, and created a model-filtered version of *X* by the element-wise product *α*_*jm*_ · *X*_*j*_, *j* = 1, …, *p*. However, rather than use a model-filtered *X* from a single subject’s condition, we chose for purposes of interpretation a “canonical” *X* derived from an average of subjects for whom their task condition predictions both satisfied the criterion that 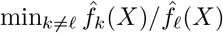 was at least 3. In our sample, 70% of the subjects satisfied this criterion. Figure 2 (a)-(i) show the main features of the model for interpreting the spatial, spectral, and temporal aspects that contribute to successful performance for the Go/NoGo task.

Figure 2(a), (d), (g) show the locations in the brain of independent components whose scalp maps fit well to scalp projections of equivalent dipoles at those locations (see also Table 1). We interpret these as the approximate locations of the independent EEG sources, given the low %RV of the fits. Figure 2 (b), (e), (h) show time-frequency maps of an augmented *X*∗ for each task condition (i.e., augmented to include its original frequency resolution). Red denotes increases, and blue denotes decreases, in log spectral power relative to the baseline period. Figure 2 (c), (f),(i) show the model-filtered *X*∗ for two of the four derived units in the hidden layer, one whose contribution to 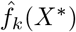 is solely due to the Go condition and one whose contribution to 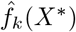 is solely due to the NoGo condition, shown in green and red, respectively. These are 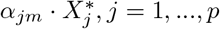. We have displayed these model “signatures” separately by component, and we have reshaped the 2-D time-frequency panel as a single 1-D profile, where the .8-second time windows for each frequency interval have been concatenated along the horizontal axis.

**Figure 2:**
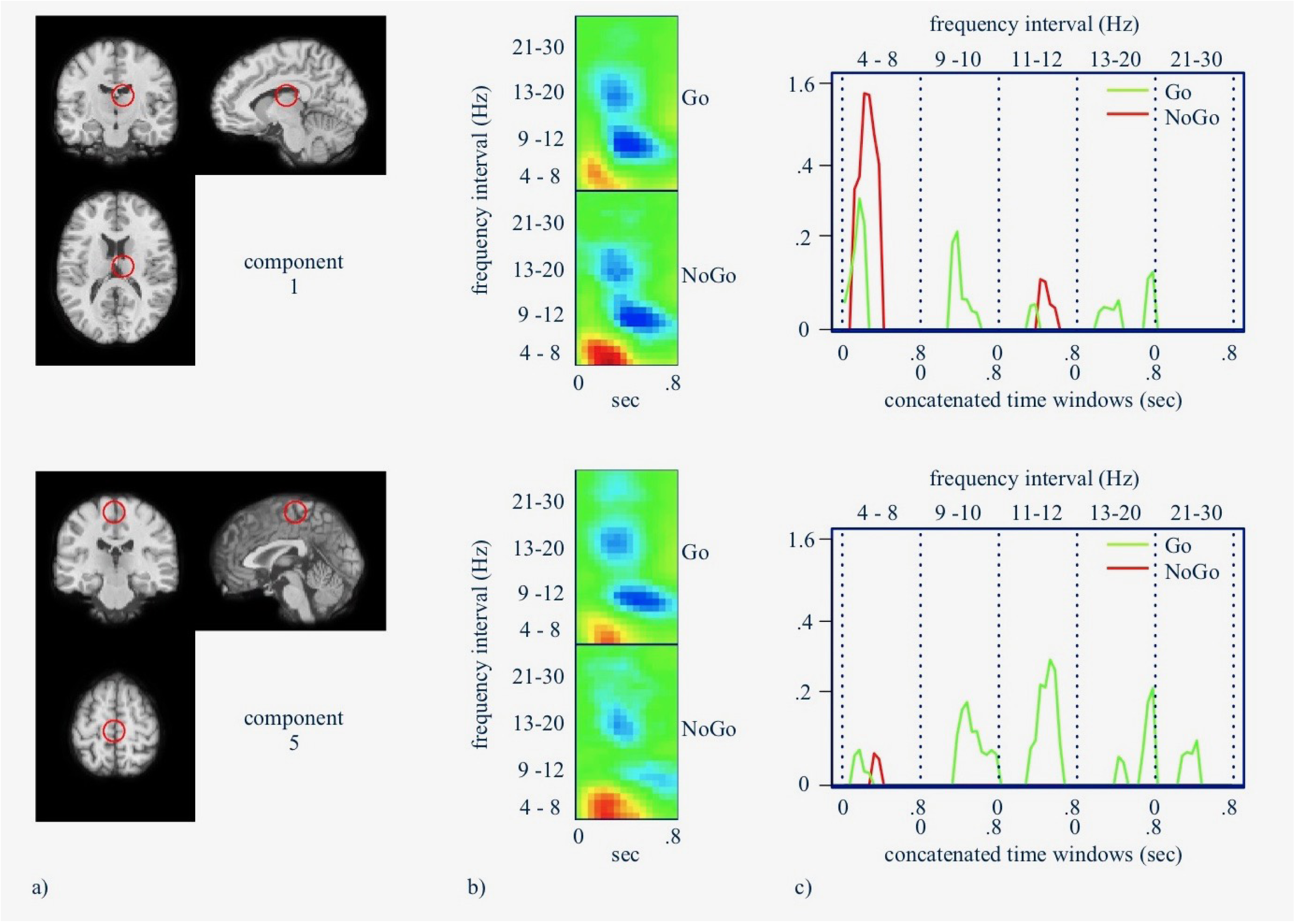
a) Approximate location of the component source having the best fit of its corresponding scalp map to the scalp projection of a single equivalent dipole. b) Log power spectra of the independent component for the .8-second time window post-stimulus. Red denotes increased and blue denotes decreased spectral power relative to the pre-stimulus period. c) Model-filtered spectral power for frequency intervals and temporal epochs that maximize the prediction functions for each *C*_*k*_(*X*). These component ‘signatures’ are viewed here as 1-D profiles with the .8-second time windows for each frequency interval concatenated along the horizontal axis.

**Figure 3:**
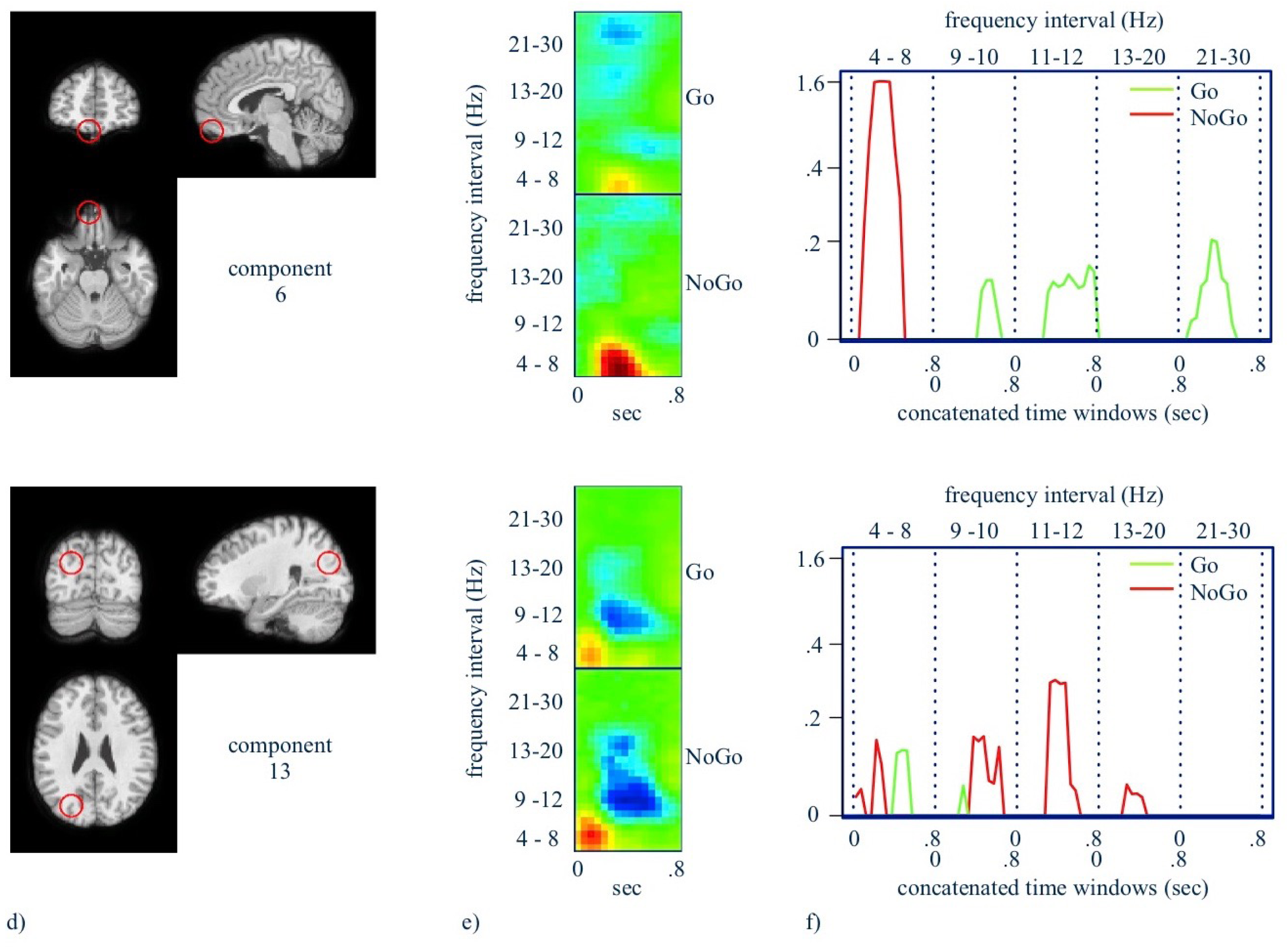
d) Approximate location of the component source. e) Log power spectra of the independent component. f) Component ‘signatures’ for prediction of *C*_*k*_(*X*). See Figure 2a for detailed description.

**Figure 4:**
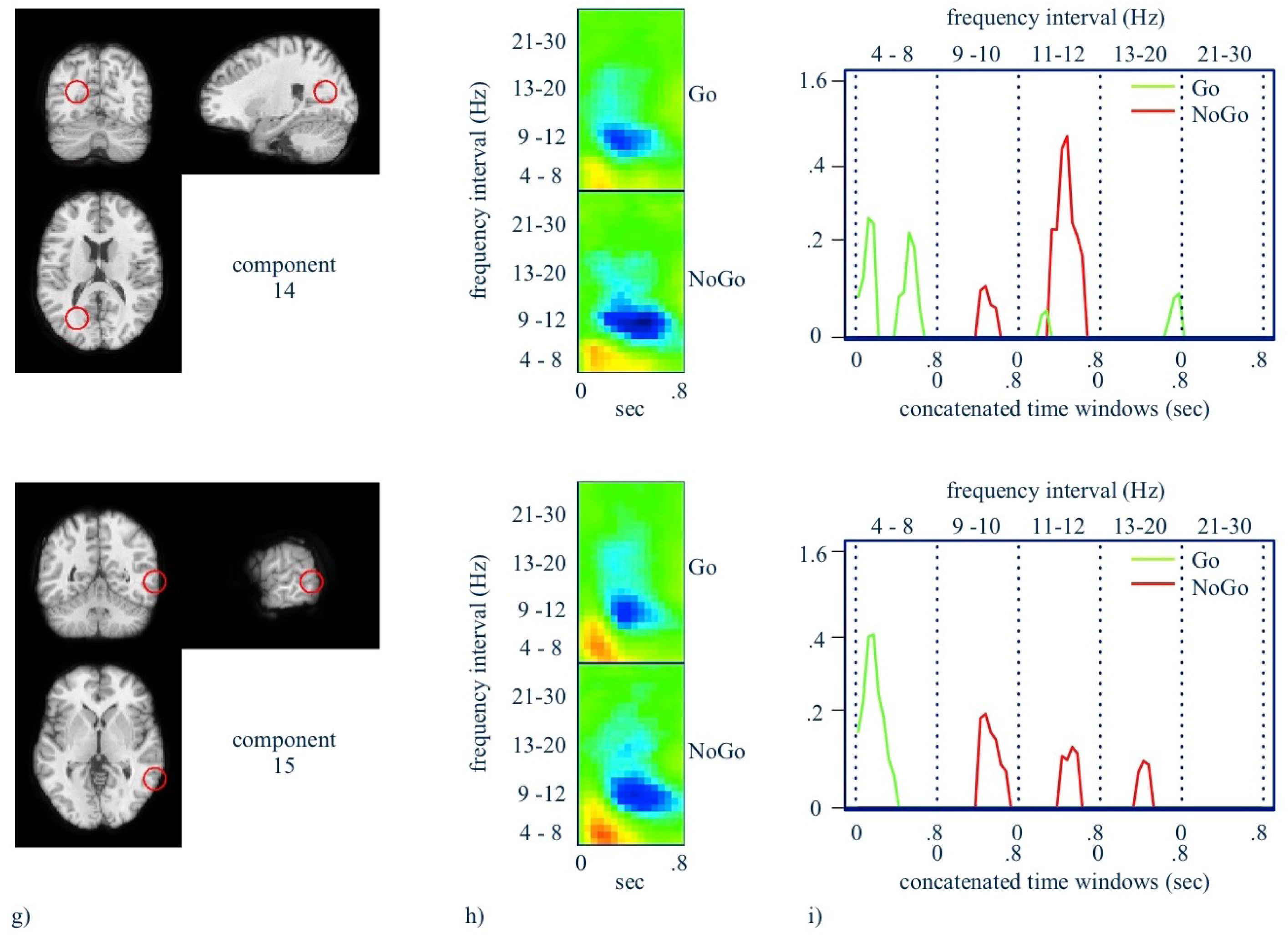
g) Approximate location of the component source. h) Log power spectra of the independent component. i) Component ‘signatures’ for prediction of *C*_*k*_(*X*). See Figure 2a for detailed description.

All model-filtered peaks are positive because important event-related increases in log spectral power have positive *α*_*jm*_, and important event-related decreases in log spectral power have negative *α*_*jm*_. However, the time-frequency panel reveals the sign of the important peaks. The peaks themselves are statistically significant based on 50,000 nonparametric bootstrap samples of model signatures and a Bonferroni threshold given by single-test levels of .05/(2*p*). Notable signatures for NoGo and Go trials are (see also Table 3 for full list):

NoGo trials 1. mid time course theta increases in the thalamus, left preSMA, left orbitofrontal cortex, and left superior parietal cortex/precuneus;
2. mid time course with upper alpha decreases in the thalamus, left superior parietal/precuneus, left occipital/cuneus, and right middle temporal gyrus;
3. mid time course lower beta decreases in the left superior parietal/precuneus and right middle temporal gyrus;
Go trials 1. early theta increases in the thalamus, left preSMA, left occipital/cuneus, right middle temporal gyrus, mid time course lower alpha decreases in the thalamus, left peSMA, left orbitofrontal cortex,
2. mid time course upper alpha decreases in the thalamus, left orbitofrontal gyrus,
3. late lower beta increases in the thalamus, left preSMA, and left occipital lobe/cuneus,
4. mid time course upper beta decreases in the left preSMA, left orbitofrontal cortex.

**Table 3:**
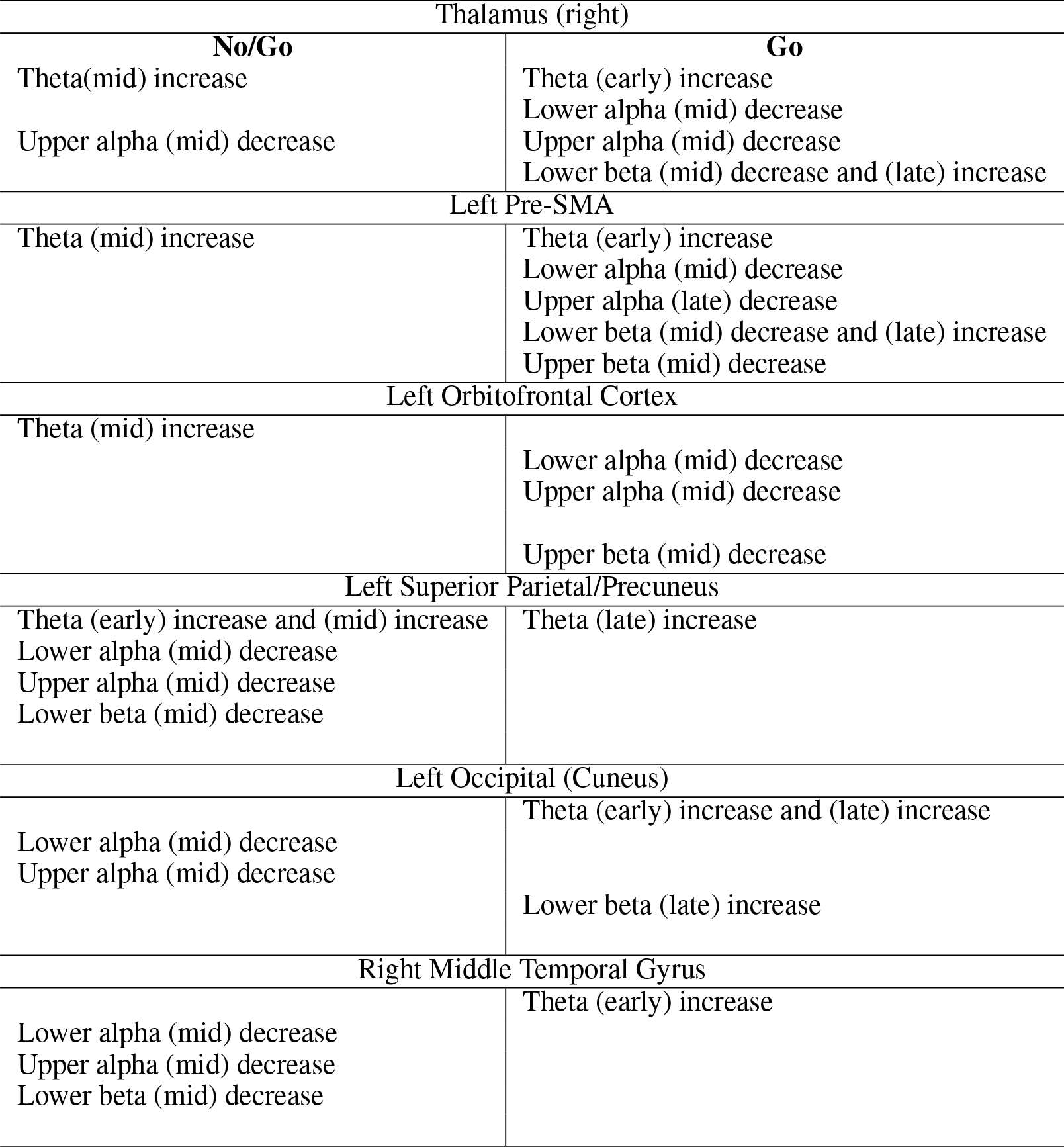
Listed are the 6 components that delineate Go and NoGo trials designated by region with the EEG time frequency (theta, lower alpha, upper alpha, lower beta, upper beta) and relative time (early, mid, late) of their peak in the stream of processing for each type of trial.

## 4 DISCUSSION

In this study, we used a machine learning approach to identify ERSPs that discriminate between selection and inhibition in a conventional Go/NoGo task. Our modeling provides a detailed account of time-frequency patterns that differentiate Go and NoGo trials. The detailed accounting of EEG time-frequency patterns localized to brain sources allows for elaboration on previous findings from fMRI and EEG studies by providing more information about EEG power changes in multiple frequency bands per region underlying the selection and inhibition processes engaged in the Go and NoGo trials, respectively. Additionally, the modeling revealed multiple relevant, localized time-frequency changes not previously discussed in earlier selection and inhibition models.

Through our modeling, we identified six components from which neural signals across the EEG spectra, ranging from theta to beta, contributed uniquely to selection and inhibition. Mid-trial processing in inhibition involved theta increases in thalamus, preSMA, frontal pole and left parietal/precuneus cortex. Mid-trial processing in inhibition also involved 1) decreased upper alpha in thalamus, left parietal/precuneus region, left occipital/cuneus, and right middle temporal gyrus, 2) decreased lower alpha in left parietal/precuneus region, left occipital/cuneus, right middle temporal gyrus, and 3) decreased lower beta in the left parietal/precuneus region and right middle temporal gyrus. Early processing in selection involved 1) theta increases in thalamus, preSMA, left occipital/cuneus region, and right middle temporal gyrus. In selection, mid-trial processing involved 1) decreases in lower alpha in thalamus, preSMA, and frontal pole, 2) decreases in upper alpha in thalamus and frontal pole, 3) decreases in lower beta in the thalamus and preSMA, and 4) decreases in upper beta in preSMA, and frontal pole. Finally, in selections, late processing involved 1) theta increases in left parietal/precuneus region, left occipital/cuneus region, 2) upper alpha decrease in preSMA, 3) lower beta increases in thalamus, preSMA, and left occipital/cuneus. These distinctions between Go and NoGo EEG patterns from these sources were shown to be reliable by 5-fold cross-validation, yielding highly accurate prediction rates.

Our findings are consistent with previous findings showing that medial frontal areas play a role in both response selection and inhibition [25]. In fMRI, pre-SMA and frontal-pole (components 5 6 in the present modeling) has shown selection- and inhibition-related activation but greater inhibition-related activation [16, 1]. In other studies, the pre-SMA showed greater than baseline fMRI signal for both Go and NoGo trials. A meta-analysis showed that several regions constitute a cognitive control network essential to performing the Go-NoGo task, with the following regions associated with specific cognitive operations, with the parietal lobe with the decision to act or not act in Go-NoGo conditions and subsequently attenuating activity in preSMA and motor cortex for NoGos, and preSMA with inhibiting motor responses and conflict detection, adjusting response thresholds, and switching from one action plan to another [1].

Our results suggest greater neural synchrony in the theta band in inhibition versus selection and possibly reduced alpha synchronization for selection compared to inhibition, underlying the noted activation differences in fMRI. Previous time-frequency studies of the Go/NoGo task showed EEG power changes in the theta and alpha bands, with theta-band increases during the NoGo trials, and to a lesser degree during the Go trials, over midline frontal EEG electrodes [12]. Our modeling further characterizes this theta-band EEG finding and delineates early increases in the preSMA for Go trials relative to NoGo trials and early increases sustained over the trial in frontal-pole for NoGo trials. The temporal and amplitude differences in preSMA might reflect a bias toward selection, possibly related to early template matching on Go trials [26, 27] and added matching-related processing when infrequent NoGo trials occur. The theta power increase in frontal-pole is consistent with this region’s involvement in inhibitory control [1].

Our previous EEG time-frequency study also demonstrated declines in alpha power (i.e., desynchronization of regional neural oscillations from baseline) for both Go and NoGo stimuli in midline regions [12], with the distribution across both stimulus types from this model implicating a common cognitive mechanism across tasks, possibly related to evaluation of the stimulus properties. The present modeling localized corresponding midline alpha power changes to preSMA and frontal pole. The model, however, shows that the observed alpha desynchronization within these regions was associated with selection and not inhibition. Additionally, however, the modeling reveals inhibition-related alpha desynchronization within the posterior visual/visual association areas that occurred later in the trial. Alpha-band oscillations have been imputed to represent inhibitory and timing processes linked to attention suppression and selection [28]. Across various processing domains, alpha power has been shown to increase in systems targeted for disengagement from a processing task (e.g., the dorsal visual stream when ventral visual stream processing is required [29]) and decrease in systems targeted for engagement in the task. Within the alpha band, desynchronization or power reductions from baseline are imputed to index a release of “alpha” suppression; whereas synchronization or power increases are imputed to index an increase in “alpha” suppression. Further, desynchronization of lower alpha has been associated with attention allocation and desynchronization of upper alpha has been associated with semantic retrieval processes [30]. Thus, the mid-to-late increases in lower and upper alpha desynchronization within posterior visual/visual-association regions on NoGo trials might reflect additional attention and semantic processing taking place after earlier inhibitory control signals from preSMA and frontal-pole were initiated. An alternative general account for alpha attenuation in posterior cortical regions is that when midline theta (and likely beta) activation is present, posterior cortical regions encoding representations of items will be characterized by suppressed alpha [31, 32]. This would be be more pronounced for the NoGo trials where disengagement of posterior regions would be indicated if the identified stimulus differs from the predominantly present target.

The Go responses also differ from the NoGo responses with the presence of lower and upper beta-band EEG power changes in the processing stream. The late beta EEG power change is likely associated with final object identification for a decision in the Go trials, and is detected as late beta increases in the model at the preSMA, thalamus and the left occipital region, which are all part of the object identification and name retrieval network [33, 34]. This network for Go trials is in keeping with the need to clearly designate an item as a Go target by identifying it in semantic memory, which has been identified previously to occur via a preSMA-thalamic-cortical representation circuit via beta rhythms [34]. The network linked by early theta-band EEG power changes in the preSMA and posterior cortical regions during Go trials are consistent with what has been found in previous studies, with frontal theta being sensitive to increasing amounts of information prior to and upon identification of the stimulus for a Go decision. Such theta EEG signals may reflect communications between posterior cortical regions involved in visual stimulus processing and object identification as noted aboves. The alpha power changes in the preSMA, orbitofrontal and thalamic regions are proposed to be associated with anticipatory rule updating, motor response [35], and or item search.

Finally, our model may have utility in the context of recent advances in non-pharmacological therapeutics in neurological illnesses, chiefly, the use of electromodulation in the treatment of neural dysfunction. A key parameter of neuromodulation is the frequency of the stimulation if alternating current/magnetic fields are involved. Thus, models based on EEG frequency may lead to better insight not only to the dysfunctional circuitry underlying a patient’s condition, but also providing insights into the type of electromodulation that may be effective in remediating the dysfunctional state. In addition, time-frequency models allow for making predictions and generating hypotheses concerning nodes and frequencies in the model that would be proposed to change with application of electromodulatory stimulation at specific sites. Electrically manipulating the nodes or connections in the brain associated with regions in the model while monitoring accuracy and reaction time will help in refining the model.

## 5 ACKNOWLEDGEMENTS

We would like to thank Scott Shakal, Rachel O’Hair, and Kylee Yeatman for editorial assistance on this manuscript and cleaning the EEG data.

## 6 FUNDING

The authors received no specific funding for this article.

